# Measuring Joint Pain Through Tibio-Femoral Flexion: technique validation and assessment of behavior in response to different analgesics in rats

**DOI:** 10.1101/2025.08.13.669963

**Authors:** Kauê Franco Malange, Douglas Menezes de Souza, Júlia Borges Paes Lemes, Luciane Candeloro Portugal, Carlos Amílcar Parada

**Author notes:** ^¥^These authors contributed equally and share the first authorship of the manuscript (Kauê Franco Malange, Douglas Menezes de Souza).

## Abstract

**Background:** Assessing knee joint pain in experimental OA (EOA) models remains a significant challenge. Our study demonstrates that the use of an adapted electronic von Frey (aVF) device, featuring a modified tip, surpasses the standard von Frey (sVF) in detecting knee joint pain behavior and evaluating the efficacy of analgesic treatments in OA model induced by monoiodine acetate (MIA).

**Results:** The sVF was able to induce a behavior profile in naïve animals characterized by a hind paw flinching and withdrawal reflex. This behavioral response was affected by intraplantar lidocaine, which prone to the increase in mechanical thresholds related to sVF and validate it as pain related behavior. On the aVF method, the animals displayed no alterations at their mechanical thresholds in presence or absence of lidocaine, suggesting minimal stimulation of hind paw by the modified methodology. In animals where an osteoarthritic phenotype was induced by MIA, the aVF was able to detect a significant reduction on joint mechanical thresholds. The behavior linked to aVF was significantly affected by systemic delivery of morphine, confirming a nociceptive-like phenotype and suggesting a pain behavior predominantly triggered by joint flexion. The aVF was also able to detect an analgesic profile in MIA-OA rats treated with dexamethasone, LPS-RS, fucoidan, and morphine, indicating effectiveness in measure different the response profile triggered by analgesic drugs that affect joint pain perception.

**Conclusions:** Our results suggest aVF as a more appropriate method to evaluate joint pain in rats.

## 1. INTRODUCTION

Osteoarthritis (OA) is a widespread and debilitating condition, primarily affecting older adults (Cui et al., 2020). Clinically, OA is characterized by joint swelling, stiffness, inflammation, and progressive degeneration, ultimately leading to chronic pain and significant physical disability (Ji & Zhang, 2019).

To evaluate joint pain in osteoarthritic animals is a challenge. Several methods have been employed to provide accurate information regarding the pain phenotype linked with joint injury and its associated dermatomes(Malfait et al. 2013b). Typically, these methodologies are intent to analyze the static or dynamic joint performance and its outcomes in the presence or absence of the insult over the time-course related to the animal model, as seen at Weight-Bearing, Catwalk or von Frey methods (Malfait et al. 2013b; Piel et al. 2014).

Although there is a significant covariance between joint injury, antalgic gait and tactile mechanical thresholds attributed to skin dermatomes in rodents, there is some inconsistency in effective translating the correct pain phenotype in arthritic models(Malfait et al. 2013b; Piel et al. 2014). This is linked to the fact that the joint phenotype in arthritic animals is dynamic, commonly associated with the depreciation of joint mechanobiology by an inflammatory input or to primed afferent terminals disposed at the skin, which can be activated by joint motion, mechanical pressure or light touch.

Establishing a reliable pain model and employing precise tools for measuring joint pain is critical for advancing research and therapeutic interventions (Abboud et al., 2021). The accuracy of behavioral tests to measure intrinsic joint pain impacts better approaches for repair of OA-Knee joints, once humans and experimental OA animals (EOA) display similar OA outcomes (Eitner et al. 2017).

Our research group has previously published a modified von Frey method to evaluate joint pain linked with tibio-tarsal joints (Guerrero et al., 2006). This method allows a better assessment of joint mechanical thresholds as is predominantly linked with joint flexion/motion rather than paw stimulation in rodents. On the other hand, it is understood that osteoarthritis has a significant incidence in tibio-femoral joints rather than other synovial joints (Cui et al. 2020).

Thus, in this work using an adapted electronic von Frey (aVF) device we aim to validate a new method to evaluate joint pain in rats. This slight modification from a standard electronic device von Frey (sVF) into an adapted one (aVF) allows a more sensitive methodology for intrinsically evaluating knee joint pain in EOA animals. The effectiveness of the new methodology in detect different analgesic profiles linked with experimental drugs was assessed in a time-course manner at the acute and late phase of MIA model, where a histopathological assay was performed in intent to corroborate the behavioral findings.

## 2. MATERIALS AND METHODS

### 2.1. Animals

Male Wistar rats weighting from 200 ± 20g were provided by the Central Vivarium of the State University of Campinas (UNICAMP) – CEMIB. The experiments were approved by the Ethics Committee from the Institute of Biology at UNICAMP (protocol 5280-1). The animals were kept in polypropylene cages at the Maintenance Vivarium of Pain Studies Laboratory at UNICAMP with a temperature of 22 ± 2°C, light/dark cycles of 12 hours (lights on at 7:00), receiving food and water ad-libitum. The behavioral tests were performed by a blind investigator in a silent room and animals were set to acclimate for at least 1 hour before the tests.

### 2.2. Drugs and injection techniques

The drugs were injected in three distinguished routes: subcutaneous, intraperitoneal, and intra-articular. The tools and equipment for each pathway used for injection, drugs, volume, and doses are described in the Supplementary Table 1. The drugs and doses injected were based on published literature.

#### 2.2.1. Subcutaneous injection

The injections were performed at the plantar region of the metatarsal pad of the right limb of the animal, using the insulin syringe (30 G needle; BD ultrafine®). Animals were gently contained and the volume injected in each set of experiments was 50μL.

#### 2.2.2. Intraperitoneal injection

To appropriately perform the injection, animals were gently contained and inclined at 45 degrees for rostral positioning of viscera at the abdomen cavity. The drugs were injected at the lower right quadrant of the animal’s abdomen, using the insulin syringe (26G needle; Hypodermic; BD®). In each set of experiment the drugs were diluted to attend an injection rate of 10 mL/kg per animal.

#### 2.2.3. Intra-articular injection

The assessment of tibio-femoral joint was performed accordingly to (Teixeira et al. 2017; Malange et al. 2024). The animals were initially anesthetized with isoflurane (3-5%; Cristália®). After anesthesia was confirm as recommended by Guedel standard (Siddiqui and Kim 2025), animals were shaved at area around the right knee joint followed by asepsis with 70% Ethanol and Povidone-iodine (1%; Riodeine; Rioquímica®) on the tibiofemoral joint region. The right joint was flexed at 45 degrees and the intra-articular (i.a) injection was made through the patellar tendon using a 26 G (13x0,45mm; BD®) needle attached to a Hamilton syringe (Hamilton Company®). In each experiment, the overall volume did not exceed 50 μL. When two drugs were injected, a minimal interval of 30 minutes was given to avoid chemical antagonism and allow proper distribution at the synovial fluid.

### 2.3. Animal models

#### 2.3.1. Experimental osteoarthritis (EOA) induced by monoiodine acetate

The EOA model used in this study was induced by the i.a. injection of MIA (2 mg; 25 μL) as previously published (Malange et al. 2024; de Souza et al. 2024). The MIA was freshly prepared and diluted in sterile NaCl 0.9% solution and injected through intra-articular route in the right knee of the animals. The behavioral experiments were conducted pre (baseline readings) and post-MIA injections following a 21-day time-course.

#### 2.3.2. Acute arthralgia induced by carrageenan

The experimental arthralgia was induced by i.a injection of carrageenan as previously published by (Teixeira et al. 2017). The carrageenan (λ-carrageenan; Sigma-Aldrich; 300 μg; 25 µL) was diluted in a sterile NaCl 0.9% solution and injected at the tibio-femoral joint following the previous procedure mentioned on **2.2.3.** The animals were assessed for joint pain behavior at pre (baseline readings) and 3 hours post carrageenan injection.

### 2.4. Assessment of Pain Behavior

#### 2.4.1. Assessment of tactile mechanical thresholds by standard electronic von Frey (sVF)

The tactile mechanical threshold assessment was perform using a standard electronic von Frey device (Insight®; Brazil) and accordingly as previously published by (Vivancos et al. 2004a). The equipment consists of a force transducer where at the distal end, a polypropylene tip of 0.5 mm is attached. In an upward progressive movement, the tip is applied in a perpendicular manner to central area located at the plantar surface from the animal hind paw until a withdrawal reflex followed by flinching behavior is noticed. The force required to induce such behavior is registered on a digital analgesimeter and referred as mechanical threshold. The procedure was performed 6 times in each animal in alternate ways, where the absolute mechanical threshold was assumed as the average of these six trials. The readings were recorded before (baseline) and post treatments.

#### 2.4.2. Assessment of mechanical threshold using a Randall – Selitto apparatus

The mechanical threshold assessment was performed using a Randal-Selitto apparatus as previously published (Randall and Selitto 1957; Teixeira et al. 2014). After the acclimation, the animal hind paw was positioned at the analgesimeter, and an increased force in gram (g) with growing magnitude (10 g/s) was continually applied on the dermal surface of the hind paw until a withdrawal reflex was detected. The force value (g) to induce such behavior was registered. This procedure was performed six times, and in alternate ways, before (baseline) and post-treatments. The mechanical threshold of each animal was obtained deducting the arithmetic mean of the closest three measures on each time-point analyzed. The maximum force (g) applied ("cut- off”) for each animal was 130 g.

#### 2.4.3. Assessment of joint flexion mechanical thresholds using an adapted electronic von Frey device (aVF)

The mechanical threshold for knee joint flexion was performed using an adapted electronic von Frey device as previously published by (Guerrero et al. 2006) with some modifications. The adapted equipment used consists of an electronic von Frey (Insight®, Brazil) containing of a force transducer where at the distal end a 4.15 mm^2^ polypropylene tip is fixed and connected to a device that reads the force applied to the hind animal paw, converting it to grams and further registered on a digital analgesimeter. The stimulus is applied to the metatarsal footpad, perpendicularly and in ascending motion until the induction of tibiofemoral joint flexion followed by paw withdrawal. The force required to induce the joint flexion followed by the paw withdrawal reflex is recorded in grams and assumed as joint flexion mechanical threshold. The procedure was performed 6 times in each animal and in alternate ways, where the absolute mechanical threshold was assumed as the average of these six trials. The readings were recorded before (baseline) and post treatments.

### 2.5. Assessment of Proprioception using Rota-rod Test

The proprioception was evaluated using Rota rod equipment (Insight®, Brazil) as previously published by (Rondon et al. 2010) with some modifications. Briefly, within 24 hours before the test, animals were trained on Rota-rod equipment. In the training trials, the animals needed to maintain proper ambulation at the rotating bar at a constant speed of 18 rpm and for 180 seconds ("cut-off" time). Three attempts were made to each animal to achieve the cut-off time mark. Animals were set to rest for 5 minutes between each attempt. Only animals able to achieve 180s at the training trials were selected for testing with the experimental drugs. On the testing day, the rotating bar was set at a constant speed of 18 rpm and the latency to fall (s) of each animal was recorded as an indicative of related ambulation of each animal tested.

### 2.6. Histopathology of the Knee Joint

At 21 days post-MIA injection, the animals were terminally anesthetized with Ketamine (300 mg/kg; 200 µL) and Xylazine (30 mg/kg;200 µL), co-injected intraperitoneally and perfused through the ascending aorta with saline followed by a cold 4% paraformaldehyde (PFA) solution containing 0.1 M phosphate buffer. After the perfusion, the right tibiofemoral joint of animals was collected and left in PFA solution (4%) for 24 hours. The joints were then transferred to a new container with a decalcifying acid solution (4.5% HCl, EDTA, and Sodium Tartrate). On the 14th day, the joints were immersed in paraffin. The joint tissue was sectioned in 5 μm and stained by Gomori Thricome method. In each group, the qualitative score was determined following the histopathological analysis criteria recommended by (Pritzker et al. 2006).

### 2.7. Statistical analysis

The analysis of the results was performed using GraphPad Prism v.8 software (GraphPad™, San Diego, USA). When the analysis involved a comparison between two means, the T-student test was performed. When the comparison involved more than two means, a one-way or two-way analysis of variance (ANOVA) was performed according to the experimental design followed. When the level of significance indicated a statistical difference between the analyzed means, the Tukey test was used to compare the different analyzed groups. The adopted level of significance in the analysis of the results was pc<c0.05.

## 3. RESULTS

### 3.1. Assessment of behavioral profile in animals tested for standard and adapted von Frey

Initially, we investigated whether sVF or aVF could have a difference in the mechanical threshold baseline of naïve animals. When the standard method was performed, there was a significant increase in mechanical thresholds between the lidocaine groups compared to the naïve and saline groups (Figure 1B). This observation suggests that the original tip (0.5 mm²; Figure A-I) used on sVF lead to a gradual pressure at the soft tissue from animal hind paw, of which the behavioral outcome is a nociceptive reflex linked with the mechanical nociception. On the other hand, the slight modification performed on the tip (4.15 mm²; Figure 1A-II and A-III) significantly impacted the behavioral readings. As opposed to what was observed with sVF, when the adapted method was applied (aVF), there was no difference at the mechanical threshold readings between the groups, suggesting a minimal influence of aVF in nociception linked with hind paw stimulation (Figure 1B).

**Figure 1.**
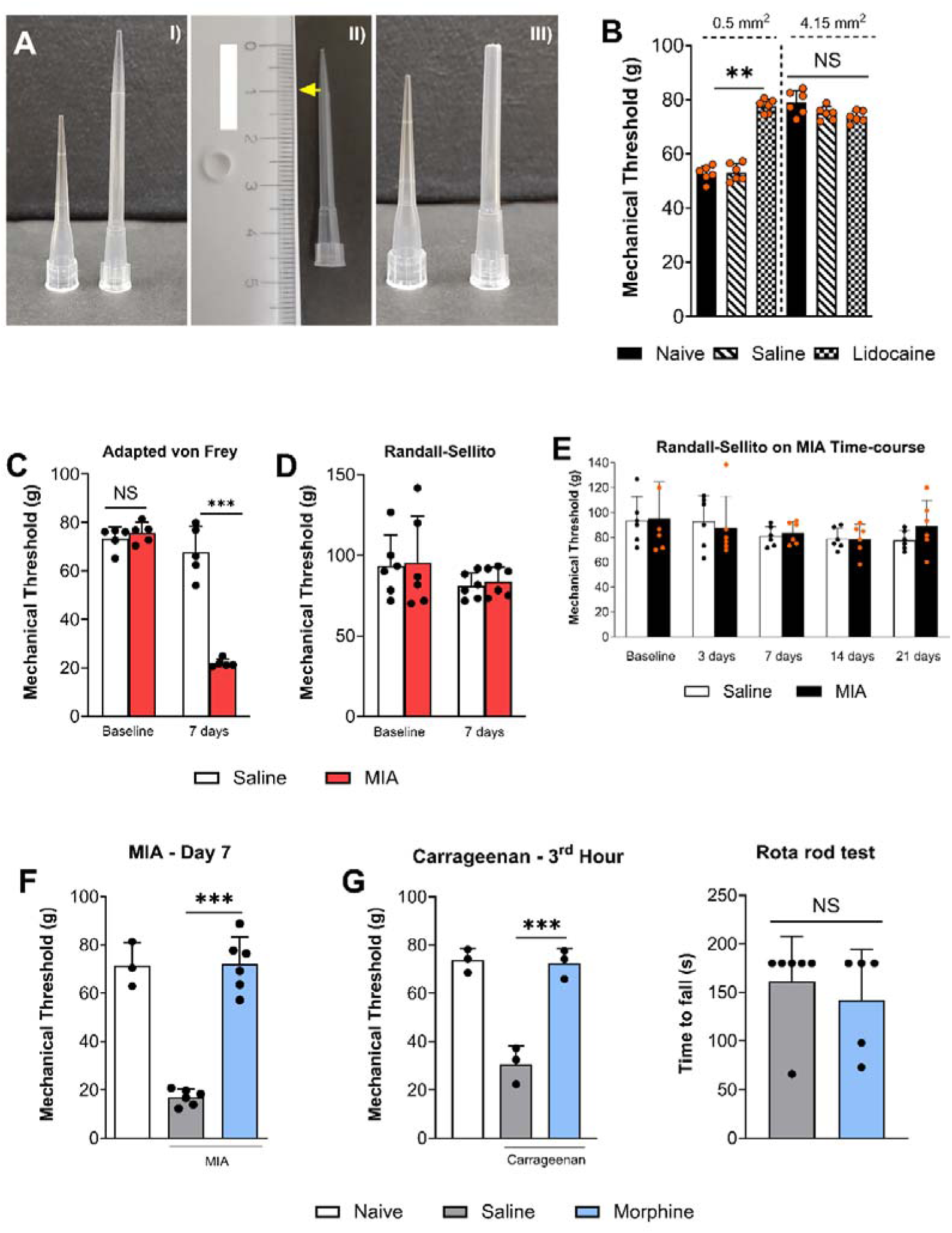
Behavioral validation of joint flexion to evaluate knee joint hyperalgesia in pain models. In (A-I to III) step by step generation of 4.15mm^2^ was used in the joint flexion test. In (B) assessment of behavior profile in naïve, saline (50 μL; intra-plantar) or 2% lidocaine (50 μL; intra-plantar) treated animals evaluated using the electronic von Frey coupled to pipette tips with different sizes. In (C) and (D), assessment of behavior profile on baseline and 7 days after MIA (2 mg; 25 μL; intra-articular) insult using the joint flexion (adapted von Frey) or RS test. In (E), (F), behavioral validation of joint flexion as nociceptive-like behavior on animals injected with MIA (2 mg; 25 μL; intra-articular) or carrageenan (300 μg; 25 μL; intra-articular) and post-treated with saline (10 mL/kg; intraperitoneal) or morphine (8 mg/kg; 200 μL intraperitoneal). In (G), motor function assessment is performed using the morphine dose in the behavioral tests. Results are shown as Mean ± Standard Deviation. Symbols ** and *** indicate p<0.01, and p<0.0001 for comparison between highlighted groups. On (B), (E), and (F), One-ANOVA with Tukey post-hoc test. On (C) and (D), Two-way ANOVA with Tukey post-hoc test.

### 3.2. Assessment of knee joint pain behavior using the adapted von Frey

After establishing a routine baseline reading for the aVF, our next experiment was to pursue if the new method was able to assess joint pain behavior in animals conditioned with different sets of stimuli that typically lead to a sensitize state in rodents, such as MIA and carrageenan (Schuelert and McDougall 2009; Teixeira et al. 2017). In our hands, when animals were previously inoculated with MIA or carrageenan and tested with aVF, we observed a significant decrease in mechanical thresholds in comparison with saline or naive controls (Figure 1C, F and G). This suggests that aVF is capable of induce a knee joint flexion, an event that naturally led to the firing of joint nociceptors (Schuelert and McDougall 2009). On the other hand, when animals were tested at the Randall-Sellito (RS), that typically does not lead to flexion of the knee joint, no differences were noted at the mechanical thresholds between MIA and Saline groups (Figure 1D). In addition, on a 21-day time-course, both saline and MIA animals display similar mechanical thresholds when tested with RS method (Figure 1E). Of note, in spinal cord injury models, the mechanical thresholds linked with the neuropathic pain phenotype can be assessed using RS method in up to 72 days after injury (Santos-Nogueira et al. 2012). Lastly, the behavior linked with aVF can be assumed to be nociceptive related, as the significant decrease in mechanical thresholds seen both at MIA and carrageenan challenged animals were reversed by systemic delivery of morphine (Figure 1F and G) without impact on motor function (Figure 1H).

### 3.3. Assessing the analgesic profile of different drugs during acute joint pain induced by MIA: accuracy of adapted von Frey

After validating the aVF test for measuring knee joint pain, we aimed to investigate the effectiveness of the aVF to assess the different analgesic profile of drugs known to impact OA joint pain, as morphine, dexamethasone and fucoidan (Fernihough et al. 2004; Huebner et al. 2014; Lee et al. 2015b). We also include in our runs LPS-RS (TLR4 antagonist) given potential implications of TLR4 in joint pain and repair (Wang et al. 2020; Kim et al. 2020). In our hands, the pre-treatment with dexamethasone, LPS-RS and fucoidan significantly affected MIA acute pain in a different manner (Figure 2B-E). Initially, MIA triggered a profound decrease in mechanical thresholds, starting at the 1^st^ hour and displaying similar values up to 72 hours after stimulus (Figure 2B). The pre-treatment with dexamethasone was able to mitigate MIA-OA pain effectively at the 2^nd^ hour after stimulus (Figure 2C). For fucoidan, effectiveness was reached at the 4^th^ hour after stimulus (Figure 2E). With LPS-RS, an effective steady analgesic profile was observed between the 2nd and 6^th^ hour after stimulus (Figure 2D). None of these analgesic drugs was able to display residual effects at 24 hours after MIA stimulus.

**Figure 2.**
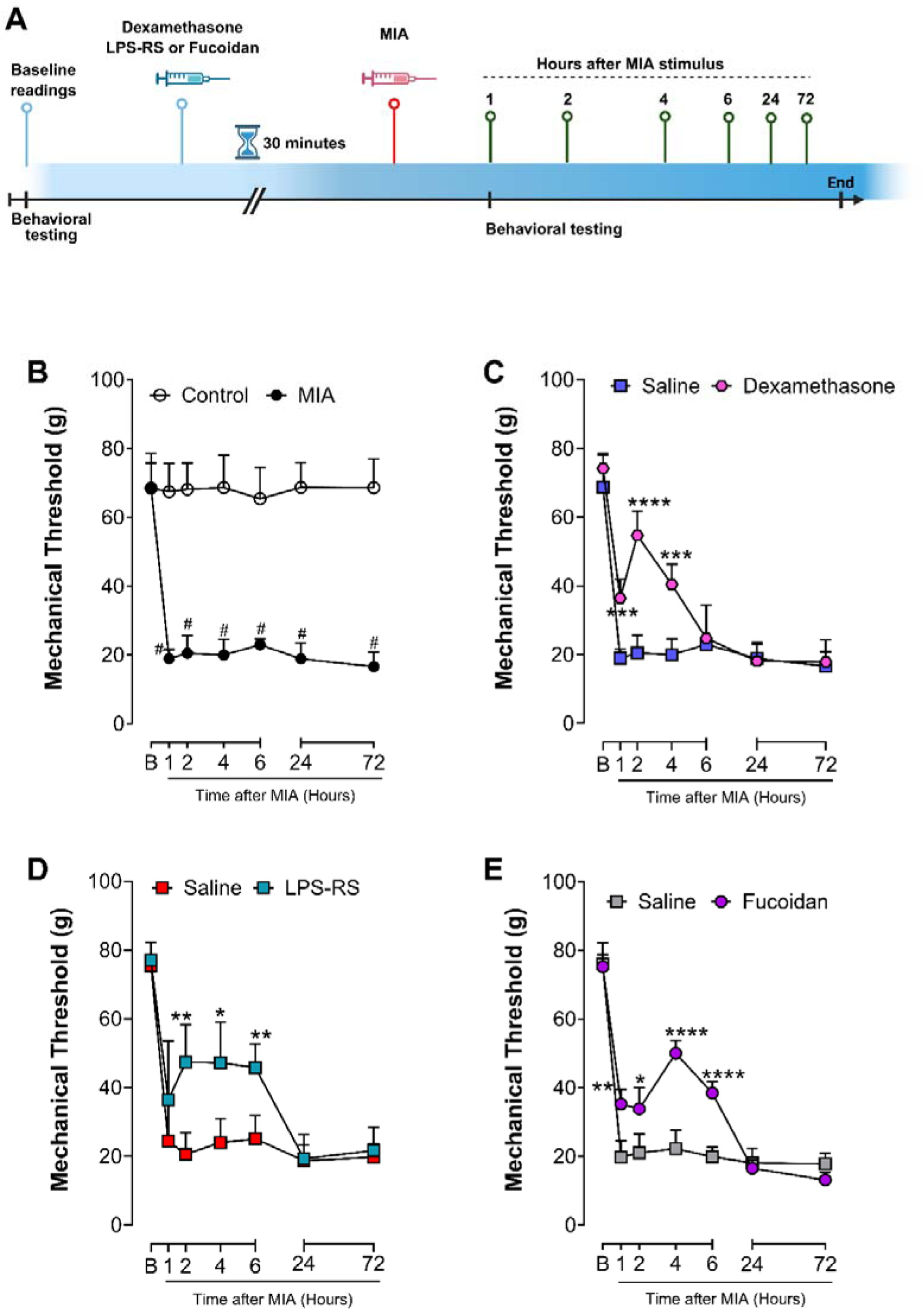
Assessment of effectiveness from adapted von Frey on acute joint induced by MIA. From (B-E) The acute joint pain evoked by MIA (2 mg; 25 μL; intra-articular) is significantly reversed in animals’ pre-treatment with dexamethasone (12 µg/µL; 25 µL; intra-articular), LPS-RS (1 µg/µL; 25 µL; intra-articular) or Fucoidan (50 mg/kg; 200 µL; intraperitoneal) in comparison with saline controls. Results are shown as Mean ± Standard Deviation. On (B) symbol # indicates p< 0.05 for comparison with the control group. From (C-E) symbols *, **, *** indicate p<0.05, p<0.01, and p<0.0001 in comparison with Osteoarthritic + Saline control (Saline on C-E). Two-way ANOVA with Tukey posthoc test.

### 3.4. Assessing the analgesic profile of different drugs during joint pain induced by MIA: post-treatment paradigm and accuracy of adapted von Frey

Given the results obtained with the pre-treatment paradigm, we next evaluated the capability of aVF in detecting analgesic profiles in animals that were initially primed for OA pain. In such experiments we used the same drugs as mentioned previously and the time-course runs were performed at day 7 after MIA challenge. In our hands, the use of dexamethasone, LPS-RS and morphine in MIA-OA animals differently impacted the mechanical thresholds. Initially, all animals primed with MIA 7 days before the tests had similar mechanical thresholds (Figure 3B-E). The treatment with dexamethasone and LPS-RS only provided a significant reversal of MIA-OA pain only in the 6^th^ hour after treatment (Figure 3B, C). On the other hand, morphine was able to induce significant analgesic effects starting at the 1^st^ hour and lasting to the 4^th^ hour after treatments (Figure 3D). Surprisingly, fucoidan had no effects on the sensitization induced by MIA (Figure 3E). The results suggest the capability of aVF in detect different analgesic profiles of drugs that significantly impact OA knee-joint pain.

**Figure 3.**
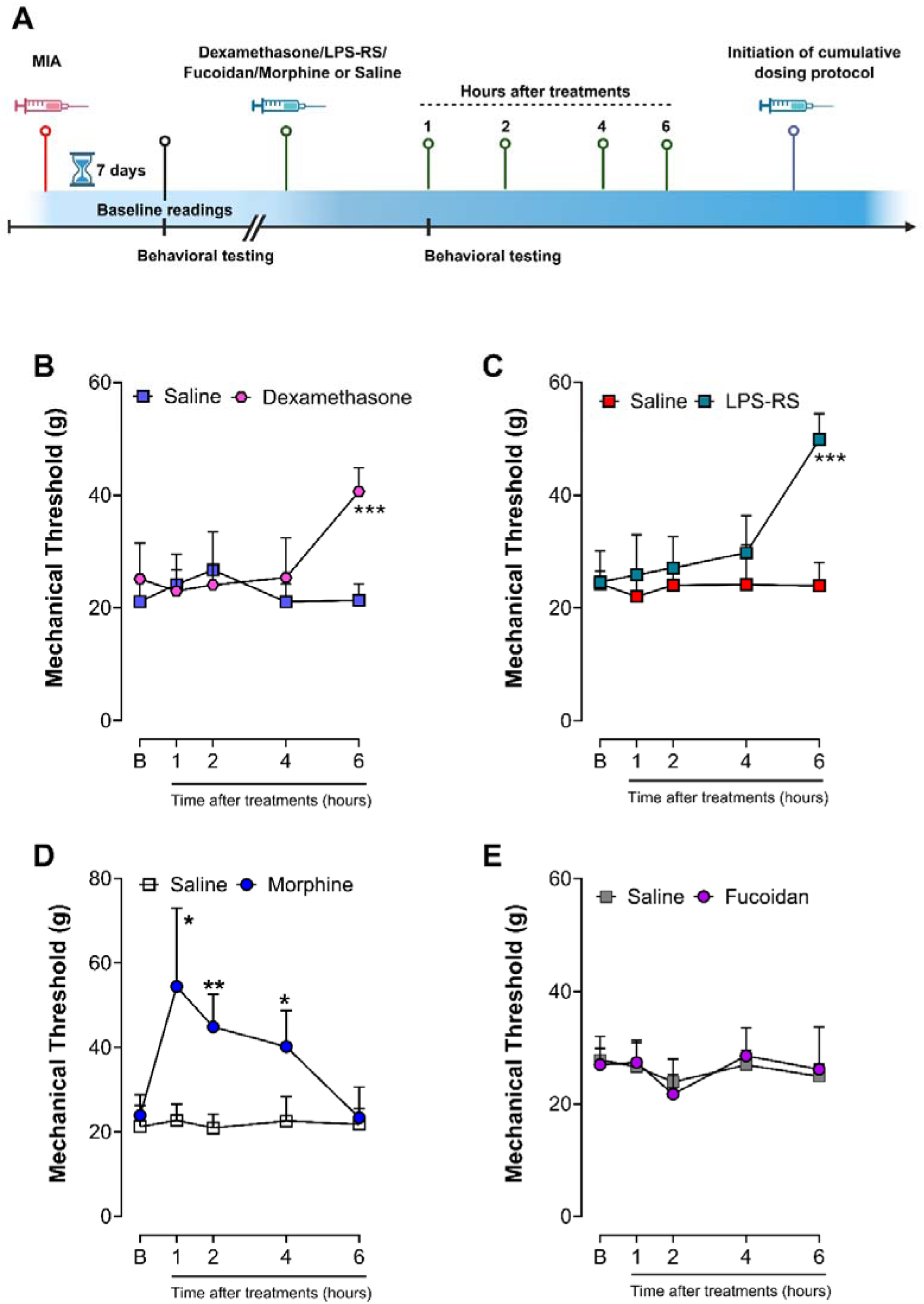
Assessing the analgesic profile of different drugs during joint pain induced by MIA: post-treatment paradigm and accuracy of adapted von Frey. From (B-E), the hyperalgesic profile of rats previously treated with MIA and tested on day 7 after insult is significantly attenuated in a time-dependent manner on animals treated with dexamethasone (12 µg/µL; 25 µL; intra-articular), LPS-RS (1 µg/µL; 25 µL; intra-articular) or morphine (8 mg/kg; 200 μL intraperitoneal), but not with Fucoidan (50 mg/kg; 200 µL; intraperitoneal). Results are shown as Mean ± Standard Deviation. Symbols *, **, *** indicate p<0.05, p<0.01, and p<0.0001 in comparison with Osteoarthritic + Saline control (Saline). Two-way ANOVA with Tukey posthoc test.

### 3.5. Assessing the analgesic profile of different drugs during joint pain induced by MIA: cumulative dosing paradigm and accuracy of adapted von Frey

As a last set of experiments intent to determine the effectiveness of aVF in measuring joint pain, we performed a cumulative dosing protocol with dexamethasone, LPS-RS, fucoidan, and morphine in animals previously challenges with MIA. In these experiments, single-dose were performed on days 7, 14, and 21 after MIA challenge with a time-course analysis being performed after the last dose on day 21. In our hands, the cumulative dosing protocol of fucoidan on days 7 and 14 was able to affect the chronic pain phenotype induced by MIA (Figure 4E). Such reversal was not seen in animals treated with dexamethasone, LPS-RS, or morphine (Figure 4B, C, and D). At the time-course analysis performed on day 21, only morphine attenuated the sensitized seen in MIA-OA animals, providing a significant increase in mechanical thresholds at the 1^st^ and up to the 6^th^ hour after treatments (Figure 4D).

**Figure 4.**
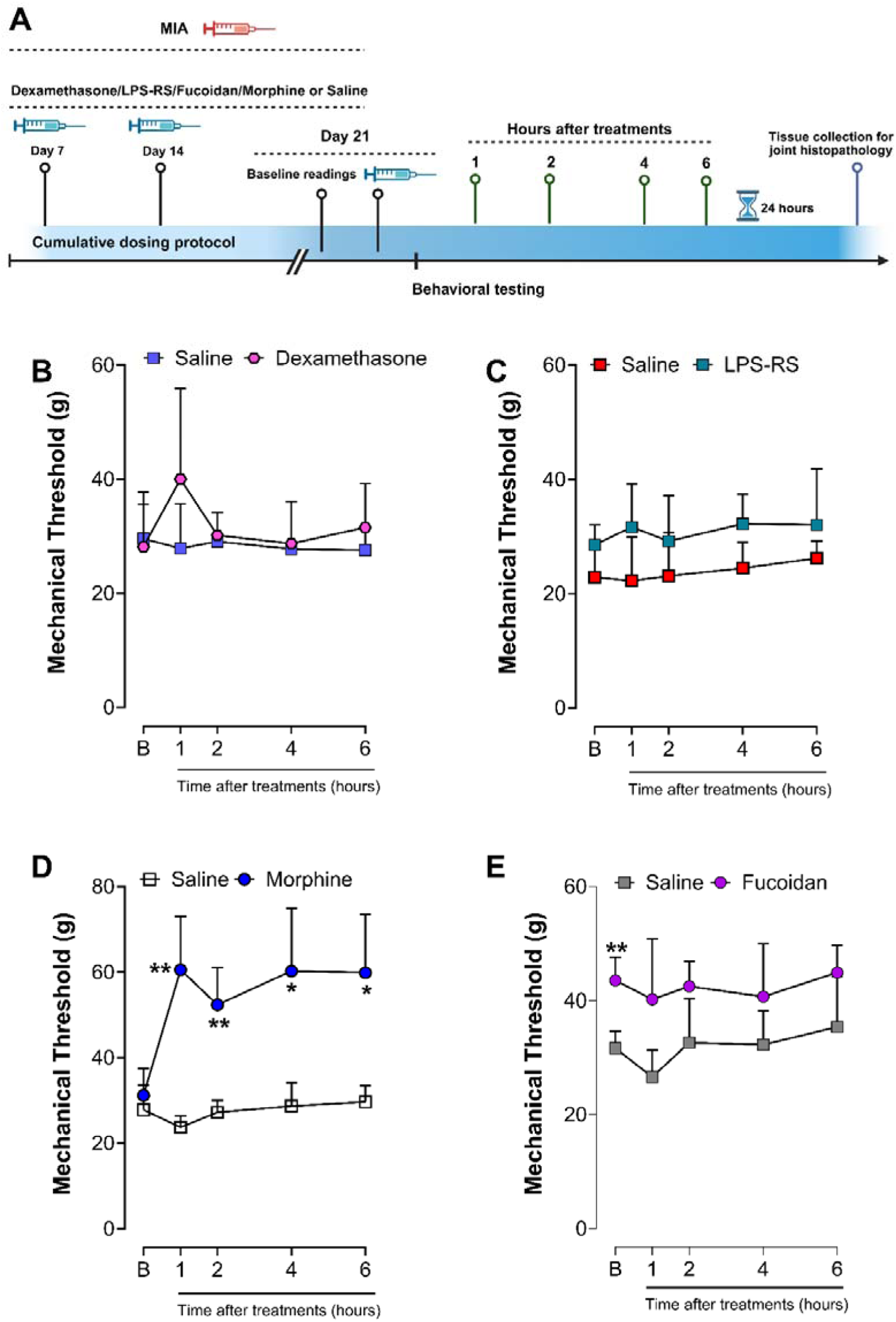
Assessing the analgesic profile of different drugs during joint pain induced by MIA: cumulative dosing paradigm and accuracy of adapted von Frey. The cumulative dosing protocol of Fucoidan (E; 50 mg/kg; 200 µL; intraperitoneal) on days 7 and 14 was able to affect the chronic hyperalgesic profile induced by MIA (P<0.01 vs OA-saline controls at baseline time-point). This reversal in chronic pain phenotype was not seen in rats treated with dexamethasone (B; 12 µg/µL; 25 µL; intra-articular), LPS-RS (C; 1 µg/µL; 25 µL; intra-articular), or morphine (D; 8 mg/kg; 200 μL intraperitoneal) ) on day 7, and day 14 after MIA insult. However, on a single-dose time-course analysis at day 21 time-point, only morphine attenuated this hyperalgesic profile induced by MIA. Results are shown as Mean ± Standard Deviation. Symbols *, **, *** indicate p<0.05, p<0.01, and p<0.0001 in comparison with Osteoarthritic + Saline control (Saline). Two-way ANOVA with Tukey posthoc test.

### 3.6. Assessment of Histopathology in animals submitted to the cumulative protocol

As an extra set of experiments, we performed a histopathological analysis in the animals submitted to the cumulative protocol with dexamethasone, LPS-RS, fucoidan and morphine to evaluate the impact of the cumulative treatments on joint morphology. For this purpose, at the end of the time-course analysis on the 21^st^-day post-MIA injection, the animals were euthanized to collect the tibiofemoral joints.

Our histopathological evaluation has shown that MIA-OA animals that were treated cumulatively with saline (OA control) display a significant loss of articular cartilage, with regions of fibrocartilage deposition and fibrosis in the bone marrow (Figure 5AI-III). We also observed that the cumulative treatment of dexamethasone (Figure 5BI-III) or LPS-RS (Figure 5CI-III) were ineffective in mitigating the knee joint degeneration. A loss of the superficial and middle zone of the cartilaginous tissue, fibrillogenesis and fissures in the cartilaginous matrix, and cell death in the middle zone of the cartilaginous tissue were detected in these groups. In animals that were submitted to a cumulative treatment with morphine, a focal loss of cartilaginous tissue with bone deformation and the presence of fibrocartilage in the medullary region of the subchondral bone were observed (Figure 5DI-III). Surprisingly, the cumulative treatment with fucoidan was able to mitigate the tissue damage in comparison with OA controls, minimizing the loss of articular cartilage, focal areas of fibrillogenesis in the cartilaginous matrix, and absence of clusters, and an intact morphology (Figure 5EI– III). Overall, the assessment of histological score in all groups revealed that fucoidan was effective in reducing histopathological score compared to OA controls and morphine (Figure 5F). In contrast, the other treatments (dexamethasone or LPS-RS) have no impact on histopathological score, displaying similar values as OA controls (Figure 5F).

**Figure 5.**
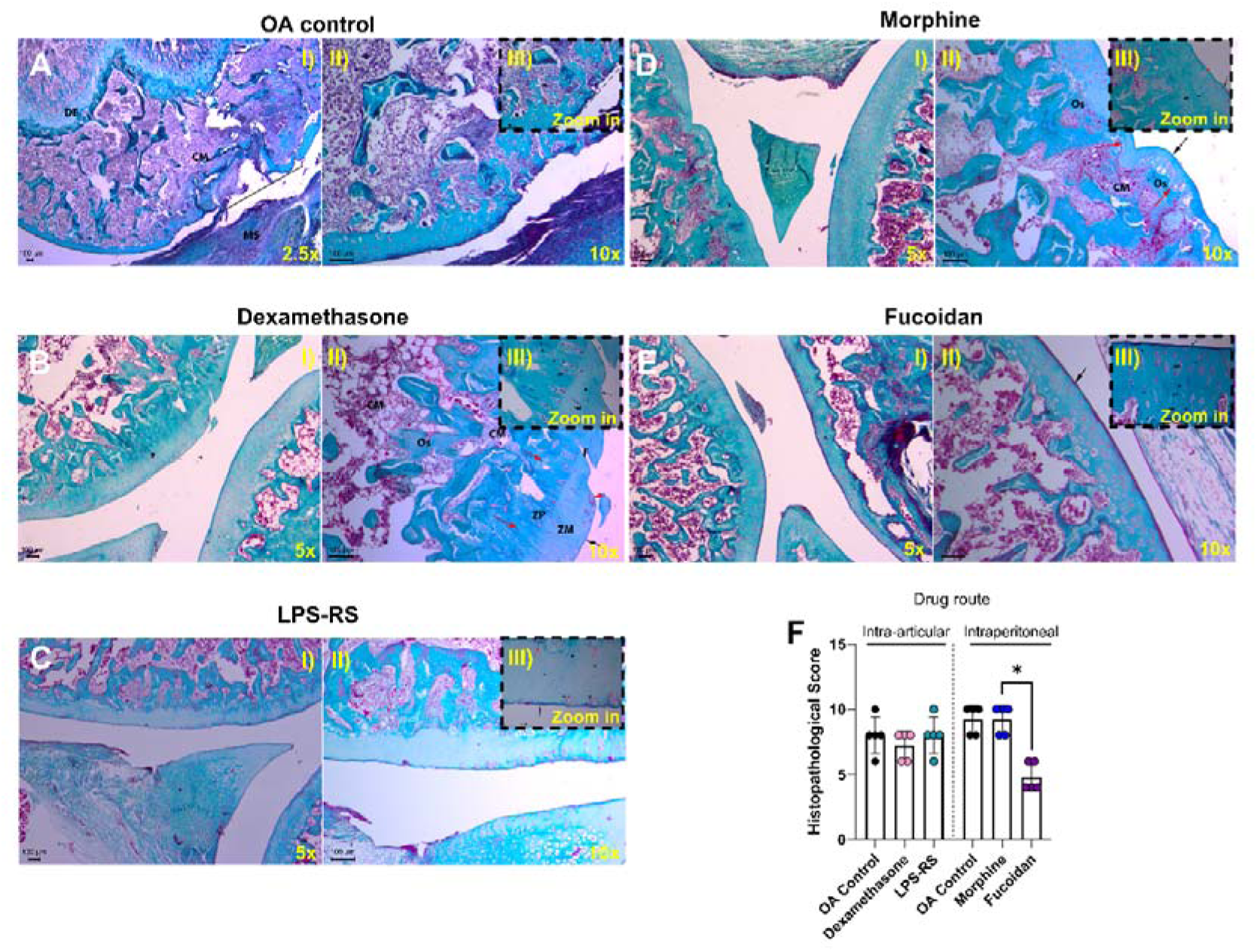
Histopathological assessment in EOA animals pre-treated with dexamethasone, LPS-RS, Morphine or Fucoidan. On (From A-E, and I-III) Gomori-trichrome stain of tibiofemoral joints from EOA controls ((a) OA control) and animals treated (b) dexamethasone (12 µg/µL; 25 µL; intra-articular), (c) LPS-RS (1 µg/µL; 25 µL; intra-articular), (d) morphine (8 mg/kg; 200 μL intraperitoneal) or (e) fucoidan (50 mg/kg; 200 µL; intraperitoneal) on 21^st^ post-MIA injection. The scale bar is displayed on the bottom left while magnification is at the bottom right. CM= medullar channel; MS=synovial membrane; OS=bone; F=cartilage fragmentation; ZP= cartilage deep zone; ZM=cartilage middle zone; Red arrow: cartilage tide mark; Black arrow: cartilage superficial zone; Asterisk= fibrilogenesis; C=cartilage erosion; Red arrowhead: chondroblasts cluster; Black line=bone remodeling. On (F), a histopathological score was performed at the OA control, dexamethasone, LPS-RS, morphine, or fucoidan groups. Results are shown as Mean ± Standard Deviation. Kruskal-Wallis test with Dunn’s posthoc test.

## 4. DISCUSSION

Evaluating animal pain in OA animal models presents a significant challenge due to pain subjective and multidimensional nature, encompassing sensory and affective components (Abboud et al. 2021). Unlike humans, animals cannot verbally express discomfort, necessitating indirect behavioral, physiological, and neurochemical assessments to infer pain levels (Miller and Malfait 2017). The traditional pain evaluation methods, such as weight-bearing distribution, gait analysis, and mechanical or thermal nociceptive thresholds, may not fully capture the spontaneous joint sensitization and functional impairments characteristic of OA (Deuis et al. 2017). On that aspect, variations in species, strain, and experimental protocols contribute to discrepancies in pain assessment, making it challenging to establish standardized and reproducible outcomes.

Assessing knee joint hyperalgesia in OA animal models is particularly complex due to the anatomical and functional characteristics of the joint, as well as the limitations of current pain measurement techniques (Malfait et al., 2013; Rios et al., 2022). Unlike cutaneous sensitization, that can be evaluated using mechanical or thermal stimuli applied to the skin, deep tissue pain in the knee joint is more challenging to quantify (von Ziegler et al. 2021). Multiple sensory fibers innervate the joint, including nociceptors that respond to mechanical stress, inflammation, and biochemical changes associated with OA progression (da Silva Serra et al. 2016). Conventional methods, such as von Frey filaments or pressure tests, often lack specificity for knee joint pain, what limit the investigator observations as secondary or referred to the primary tissue injury site.

Thus, in this work we proposed a new refined method allowing more concise observations regarding the pre-clinical evaluation of joint pain in rats. In modifying a standard electronic von Frey method, we developed a more refined method to measure knee joint pain. Initially, we noticed that naïve animals when submitted to the standard electronic von Frey have an increase in their mechanical thresholds when lidocaine was applied before testing, indicating that the mechanical pressure evoked by the tip contact with animal skin dermatomes, lead to a nociceptive reflex. On the other hand, lidocaine is unable to induce changes in mechanical thresholds in animals that were tested with the adapted von Frey methodology, suggesting a minimal interference of the adapted tip with the animal hind paw. Interestingly, we noticed a similar outcome in our previous work that has used the same method to investigate sensitization related to tibio-tarsal joint (Guerrero et al. 2006).

Joint nociceptors are stimulated mainly by harmful joint movement, presenting lower firing thresholds during an inflammation (Schuelert and McDougall 2008, 2009, 2012). At MIA-OA model, it is well established that a single injection of MIA can lead to a prominent joint inflammatory phenotype on 7 days after the insult, an event that covaries with an increase in joint pain in rodents (Schuelert and McDougall 2009; de Sousa Valente 2019). In our hands, we observed a significant reduction in mechanical thresholds on MIA-OA animals tested with the adapted von Frey, suggesting that the new method can indeed lead to a joint flexion and promote an aversive behavior in the rat. Such behavior was significantly affected by the systemic delivery of morphine, validating our observations as nociceptive-like (Fatt et al. 2024).

At standard electronic von Frey the absence of an endpoint is based on the capability from the handheld force transducer in trigger a paw tissue deformation that would eventually lead to a nociceptive reflex, read by the flinching response (Vivancos et al. 2004b). In this sense, it is important to consider that the progressive movement made by the investigator with the transducer can eventually lead to joint motion.

To emphasize the capability from the adapted von Frey in detecting joint related thresholds rather than methods that focused primary on skin dermatomes, we tested MIA-OA animals in Randall-Selitto (RS) apparatus. As expected, no differences were observed between MIA-OA and control animals tested at up to 21 days after MIA insult. Hence, these results suggest an ineffectiveness of RS in detecting oscillations on pain mechanical thresholds in MIA evoked osteoarthritis. Interestingly, similar results were obtained in previous studies (Pomonis et al. 2005; Ferreira-Gomes et al. 2008). Such results could be an indicative from a functional property of nociceptive fibers in preferentially trigger a secondary sensitization response of afferents closest to the injury site rather than at distant locations (Latremoliere and Woolf 2009).

On that note, the collective previous works (Fernihough et al. 2004; Ferreira-Gomes et al. 2008; Thakur et al. 2012; Mousseau et al. 2018) that have shown tactile allodynia measure by von Frey hairs reinforce the participation of light-touch fibers on the MIA evoked pain phenotype. Thus, it is reasonable to suggest that the adapted von Frey allows a more refined method to measure joint pain in rats, as it prioritizes the nociceptive response primary from joint, allowing a reading of the pain behavior associated directly with the injury site.

As a method to evaluate pain behavior, it is essential for a technique to be able to detect behavioral changes affected by different experimental drugs in different pain phenotypes. The injection of carrageenan at the knee-joint lead to an acute arthritis phenotype in rats with collective works showing a marked increase in nociceptive behavior peaking at the third hour after carrageenan challenge (Tonussi and Ferreira 1992, 1999; Teixeira et al. 2014, 2017). In our hands, rats that were challenged with carrageenan and tested with the adapted von Frey three hours later show a significant decrease on their mechanical thresholds in comparison with controls. Animals that were previously challenged with carrageenan and treated systemically with morphine one-hour prior testing, display significant reversal of such primed state. Thus, these results suggest an applicability of adapted von Frey to assess pain behavior in acute arthritis models as carrageenan.

The OA model triggered by MIA in murine animals display a pain phenotype typically reported as inflammatory at the early phase and shifting to neuropathic on the late phase (de Sousa Valente 2019). After validating the adapted von Frey as method to assess joint pain behavior in rats, we guided our studies in demonstrating the effectiveness from this method in detecting different analgesic profiles from drugs that had been previously reported to mitigate OA pain as morphine, dexamethasone and fucoidan (Fernihough et al. 2004; Huebner et al. 2014; Lee et al. 2015b). On the other hand, we also sought to use these drugs to provide insights regarding the phenotype associated with the pain behavior assessed in the rats.

Initially, the MIA led to a significant decrease in joint-flexion mechanical thresholds initiating at the first hour after injection and lasting up to 72 hours.

The pre-treatment with dexamethasone, LPS-RS or fucoidan was able to mitigate MIA-OA pain. Dexamethasone was able to more effectively affect MIA-OA pain at 2 hours. For LPS-RS, a significant effect was still seen around the 6^th^ hour after the MIA challenge. On the other hand, fucoidan had a more significant effect from 4 to 6 hours. Altogether, these results suggest that MIA evoked pain acutely sets an inflammatory state at the joint compartment depending on toll-like receptor 4. On that note, the long-lasting effect of TLR4 antagonist, still seen at the 6^th^ hour in comparison with dexamethasone, suggests a prominent role of TLR4 in MIA evoked pain, possibly as key mediator of inflammatory cascades triggered by damage associated molecular patterns (DAMPS) release by cartilage tissue death (Miller et al. 2015). The ability of fucoidan to attenuate joint pain more significantly from 4 to 6 hours after MIA challenge suggests an inflammatory cell recruitment event, what has been reported in other inflammatory pain models to occur during this time frame (Cunha et al. 2008; Zarpelon et al. 2013).

Once we validated our initial observations regarding the effects of different analgesic drugs in MIA acute pain, we investigated the post-treatment effects of such drugs in animals previously challenged with MIA. As the OA pain induced by MIA involves biphasic state, displaying inflammatory and neuropathic-like phenotype, we divided our behavioral experiments into 2 steps: on 7 days and 21 days after MIA challenge. On these two timepoints, the aVF method was able to detect the analgesic response of the different drugs tested. At the time-course analysis performed 7 days after MIA challenge, the treatment with dexamethasone and LPS-RS had similar analgesic outcomes, only providing significant analgesia at 6^th^ hour after treatments. Of note, TLR4 activation is known to promote downstream MAP kinase signaling, which sustained both inflammatory and neuropathic pain phenotypes through p38MAPK activity (Li et al. 2015; Luo et al. 2018). On the other hand, dexamethasone is also known to downregulate p38 expression (Lasa et al. 2002). In this sense, the similarity in analgesic activity of both LPS-RS and dexamethasone drugs in MIA-OA pain could be linked to an overlap effect regarding p38MAPK, that is also known to influence the joint tissue injury in MIA-OA model (Brown et al. 2008). The lack of effect of fucoidan in our time-course testing suggests that blocking leucocyte joint infiltration through selectins depletion is not enough to mitigate MIA induced sensitization during the inflammatory phase, at least not in a time frame of 6 hours post-treatments. Lastly, as has been shown in other studies (Fernihough et al. 2004), morphine was able to induce significant analgesia up to 4 hours after treatments.

After performing the behavioral studies on the inflammatory phase of MIA, we investigated the effects of a cumulative dosing protocol of both dexamethasone and LPS-RS in mitigating MIA induced joint injury and pain. As a qualitative comparison, we also included morphine and fucoidan. For this study, after finishing the sets of experiments on day 7, we continued dosing the animals once a week and on day 21, a time-course analysis was conducted to analyze the effects of these different analgesics in MIA late phase.

As opposed to what was found in the behavioral studies conducted on day 7, except morphine, none of the drugs was able to induce a significant time-dependent analgesic state at the late phase. Unexpectedly, the cumulative treatment with fucoidan was able to mitigate MIA chronic sensitization and joint injury. Of note, the cumulative treatment with dexamethasone, LPS-RS or morphine had no effect on MIA joint injuries.

Fucoidan has secondary activities in addition to the inhibition of selectins, where in vitro inhibition of COX-2 expression in chondrocytes, inhibition of microglial NF-kB expression, inhibition of the synthesis of cytokines (IL-1β, TNF-α, IL-6) and inflammatory mediators (PGE2 and NO), and chemokines (MCP-1) are activities discussed(Park et al. 2011; Lee et al. 2015a; Phull and Kim 2017; Phull et al. 2017; Sudirman et al. 2018; Li et al. 2021). In our studies, it seems fair to suggest that the administration route (intraperitoneal/systemic) used in the analysis would have interfered with the immediate effect of this drug, allowing it to act first on the vascular component of the animal, inhibiting only the migration of cells to the recruited site. Thus, this would not have an immediate effect on the joint hyperalgesia previously established in the model, since all the inflammatory events responsible for sensitization in joint nociceptors continue to occur independently of leukocyte migration to the joint capsule, being guided by cells resident in the joint. However, in a cumulative treatment, such combination of biological activities could have impacted both the chronic pain phenotype and joint morphology, reflecting changes in the pain behavior at the late phase and histopathological improvements.

It is well known that in humans, OA joints display high expression of TLR4 (Park et al. 2020). In rodents, at both acute and late phase of MIA model, a single intra-articular injection of TAP-2, a TLR4 antagonist, can induce a reversal of tactile allodynia when animals were evaluated 3-5 days after the injection (Park et al. 2020). On the other hand, blocking TLR4 in joint cells sometimes does not represent an interesting pharmacological approach, since it regulates primary repair responses in chondrocytes, as autophagy and oxygen damage responses (Wang et al. 2020).

Given these dynamics in TLR4 activity, it is reasonable to suggest that the antagonism of TLR4 during the inflammatory phase of MIA-OA model is beneficial since it prevents inflammatory tissue damage resulting from the apoptotic action promoted by cytokines and free radicals such as nitric oxide. However, during the process of resolving inflammation, cell survival and anti-apoptosis mechanisms are crucial for the repair and maintenance of homeostasis in any tissue, especially in cartilage tissue, which lacks vascular supply and is extremely susceptible to injury due to oxidative stress. Thus, the continued treatment with LPS-RS would not lead to a benefit at the late phase, as it would impair the recovery of cartilage homeostasis, eventually leading to no improvements in cartilage morphology.

Furthermore, it is also important to highlight that Chrysis et al. (2005) identified that dexamethasone can promote apoptosis of chondrocytes in the proliferative stage, via activation of AKT kinase and subsequently Caspase-9. In this sense, it is worth highlighting that the processes of proliferation and differentiation of chondrocytes are essential for the synthesis of extracellular matrix and regeneration of injured cartilage tissue. Hence, this also emphasizes the idea that initially dexamethasone has a beneficial effect, but in the long term it significantly impairs cartilage repair.

## CONCLUSION

The adapted von Frey methodology proved to be effective for detecting joint pain, presenting greater sensitivity in measure nociceptive behavior in different models of arthritis and in response to different analgesics. Our findings highlight the importance of assessing joint pain at its primary site, allowing a better understanding from the dynamics of different analgesics in joint pain perception rather than secondary pain as it is routinely performed in osteoarthritis research.

## ETHICS APPROVAL AND CONSENT TO PARTICIPATE

The experiment was approved by the Ethics Committee on the Use of Animals (CEUA) for protocol 5280-1 on March 21st, 2022. All experiments were conducted and performed according to the guidelines of the National Council for Animal Experimentation Control (CONCEA).

## AVAILABILITY OF DATA AND MATERIALS

The datasets generated during the current study are available from the corresponding author upon reasonable request.

## AUTHORS’ CONTRIBUTIONS

Kauê Franco Malange: conceptualization, methodology, validation, formal analysis, investigation, writing-original draft, review & editing, visualization, and funding acquisition; Douglas Menezes de Souza: investigation, visualization, writing-original draft, review & editing; Julia Borges Paes Lemes writing-original draft and formal analysis; Luciane Candeloro Portugal: performed formal analysis and histological score; Carlos Amilcar Parada contributed in the conceptualization, resources, writing - original draft, writing - review & editing manuscript, supervision, project administration, and funding acquisition.

## ACKNOWLEDGMENTS

The authors gratefully acknowledge Luciane Candeloro Portugal for supporting, performing and developing the histopathological score.

## NOTE

This work was derived from the doctoral thesis of Kaue Franco Malange.

## SUPPLEMENTARY TABLES

**Supplementary table-1.** Drugs and doses used in the study.

| Drug | Dose/Route/Volume | Reference |
| --- | --- | --- |
| Monoiodine acetate | 2 mg/intra-articular/25µL | (Xu et al. 2020) |
| λ-Carrageenan | 300 µg/intra-articular/25µL | (Teixeira et al. 2017) |
| Dexamethasone | 300 µg/ intra-articular/25µL | (Grodzinsky et al.<br>2017) |
| LPS-RS | 25 µg/intra-articular/25 µL | (Rossato et al. 2019) |
| Fucoidan | 50 mg/kg/intraperitoneal/200 µL | (Lee et al. 2015a) |
| Morphine | 8 mg/kg/intraperitoneal/200 µL | (Guerrero et al. 2006) |

